# Metformin alleviates aging-associated cellular senescence of human adipose stem cells and derived adipocytes

**DOI:** 10.1101/2020.10.05.326546

**Authors:** Matthieu Mantecon, Laura Le Pelletier, Jennifer Gorwood, Martine Auclair, Michael Atlan, Bruno Fève, Jacqueline Capeau, Claire Lagathu, Véronique Béréziat

**Author notes:** These two authors contributed equally to this work (Claire Lagathu, Véronique Béréziat). Corresponding author: Véronique Béréziat -+33140011321.

## Abstract

Aging is associated with central fat redistribution, and insulin resistance. To identify age-related adipose features, we evaluated the senescence and adipogenic potential of adipose-derived-stemcells (ASCs) from abdominal subcutaneous fat obtained from healthy normal-weight young (<25y) or older women (>60y).

Aged-donor ASCs showed more intense features of aging (senescence, mitochondrial dysfunction, and oxidative stress) than young-donor ASCs. Oxidative stress and mitochondrial dysfunction occurred earlier in adipocytes derived from aged-donor than from young-donor ASCs, leading to insulin resistance and impaired adipogenesis.

When aged-donor ASCs were treated with metformin, senescence, oxidative stress and mitochondrial dysfunction returned to the levels observed in young-donor ASCs. Furthermore, metformin’s prevention of senescence and dysfunction during ASC proliferation restored the cells’ adipogenic capacity and insulin sensitivity. This effect was mediated by the activation of AMP-activated-protein-kinase.

We show here that targeting senescent ASCs from aged women with metformin may alleviate age-related dysfunction, insulin resistance, and impaired adipogenesis.

## INTRODUCTION

Adipose tissue is the largest lipid storage and endocrine organ in the body. Recent studies have highlighted adipose tissue’s critical role in age-related diseases and metabolic dysfunction. Aging is physiologically associated with fat redistribution and a metabolic and functional decline in adipose tissue, associated with oxidative stress, inflammation and fibrosis (Cartwright, Tchkonia et al. 2007, Kuk, Saunders et al. 2009, Cartwright, Schlauch et al. 2010, Tchkonia, Morbeck et al. 2010, Luo and Liu 2016, Stout, Justice et al. 2017). The age-related redistribution of adipose tissue is characterized by the accumulation of truncal fat, hypertrophy of visceral adipose tissue (VAT), and overall loss of subcutaneous adipose tissue (SCAT) (Oikawa, Owada et al. 2016, Park, Park et al. 2016, Mancuso and Bouchard 2019). An excess amount of VAT might reflect the paucity of SCAT during aging. Indeed, while accumulation of SCAT in the lower part of the body is considered to be a metabolic sink capable of buffering surplus energy, a decline in SCAT storage capacity might lead to (i) ectopic lipid deposition in the bone marrow, heart, liver, and muscles and (ii) an increase in cardiometabolic comorbidities (Stout, Justice et al. 2017). Accordingly, it has been suggested that appropriate SCAT plasticity and expandability guard against metabolic disorders (including insulin resistance) and that the age-related loss of these properties favours metabolic disorders (Pasarica, Xie et al. 2009, McLaughlin, Lamendola et al. 2011).

On the cellular level, several age-related changes in SCAT and VAT might contribute to adipose tissue dysfunction. Failure of SCAT is likely to result from the impaired recruitment of precursors and blunted adipogenesis. The identification of adipose-derived stem/progenitor cells in the stromal vascular fraction of adipose tissue has highlighted the importance of *de novo* adipogenesis in adipose tissue expansion (Cawthorn, Scheller et al. 2012). Adipose-derived stem cells (ASCs) are defined as plastic-adherent cells expressing specific surface antigens and that are able to differentiate into osteoblasts, adipocytes and chondroblasts *in vivo* and *in vitro* (Dominici, Le Blanc et al. 2006). They can be isolated, expanded and induced to differentiate into the above-mentioned lineages by using specific culture conditions and thus constitute a useful tool for studying age-related diseases. The abundance of adipocyte progenitors/precursors that differentiate into adipocytes is an important determinant of SCAT expandability and functionality. Adipocyte-differentiated ASCs have a crucial role in lipid handling, adipose tissue expansion, and insulin sensitivity (Palmer and Kirkland 2016). With age, the frequency of ASCs decreases (Liu, Lei et al. 2017). A reduction in the ASC proliferation rate might ultimately reflect cell senescence and might be involved in the onset of metabolic alterations (e.g. insulin resistance) observed during aging.

The age-dependent senescence of ASCs has been linked to a decrease in mitochondrial activity and an increase in levels of reactive oxygen species (ROS) (Choudhery, Badowski et al. 2014, Maredziak, Marycz et al. 2016, Liu, Lei et al. 2017). Although most of the literature data show that aging has a negative effect on osteoblasts and chondrocytes, the results differ with regard to the impact of senescence on the ASCs’ adipogenic potential. Indeed, some studies found that aging had a negative effect on adipocyte differentiation (Karagiannides, Tchkonia et al. 2001, Murphy, Dixon et al. 2002, Sepe, Tchkonia et al. 2011, Caso, McNurlan et al. 2013, Beane, Fonseca et al. 2014), whereas others found that adipocyte differentiation increased with age (de Girolamo, Lopa et al. 2009, Choudhery, Badowski et al. 2014, Maredziak, Marycz et al. 2016). Lastly, it has been suggested that targeting senescent cells in adipose tissue will enhance adipogenesis and metabolic function in old age (Xu, Palmer et al. 2015).

In the present study, we found that ASCs isolated from SCAT from aged donors displayed more senescent features (such as increased expression of cell cycle arrest proteins, including p16^INK4^ and p21^WAF1^, and senescence-associated (SA)-β-galactosidase activity) than ASCs isolated from young donors. These differences were associated with a gradual decline in the proliferation and differentiation capacity during long-term *in vitro* culture. Interestingly, we showed for the first time that ASC senescence was linked to early adipocyte mitochondrial dysfunction, oxidative stress, and cellular insulin resistance.

The biguanide drug metformin is widely used to treat diabetes but also appears to modulate a number of aging-related disorders (Barzilai, Crandall et al. 2016). Metformin exerts pleotropic effects and has a favourable influence on metabolic and cellular processes closely associated with the development of age-related conditions, such as oxidative stress, inflammation, and cellular senescence. We therefore sought to determine whether metformin could prevent cellular age-related dysfunction and rescue the impaired metabolic phenotype of ASCs obtained from aged donors. Our results show that *via* the activation of AMP-activated protein kinase (AMPK), metformin alleviated the aging-associated cellular senescence of ASCs, exerted an antioxidant effect, and improved mitochondria metabolism. Lastly, metformin rescued adipocyte differentiation and function, which might be involved in the drug’s insulin-sensitizing effect *in vivo*.

## RESULTS

### Aging induces senescence, oxidative stress, and mitochondrial dysfunctions in ASCs

In order to determine the impact of aging on ASCs, we first searched for phenotypic differences between ASCs isolated from young adults (under the age of 25) and older adults (over the age of 60) during long-term *in vitro* culture. These ASCs are respectively referred to henceforth as “young-donor” and “aged-donor” cells. We evaluated the cells’ proliferative capacities, senescence marker expression, and functional impairments at early (P3), intermediate (P7) and late (P11) passages.

At P3, aged-donor and young-donor ASCs had similar proliferative abilities and thus a similar population doubling time (PDT) (Fig. 1a-b). For the young-donor ASCs, the PDTs at P7 and P11 were only slightly longer than at P3. In contrast, the PDT for aged-donor ASCs increased markedly from P5 to P11 (Fig. 1a-b), indicating a steady increase in growth inhibition. It is important to note that cell viability did not change significantly over the culture period (data not shown). The mean percentage of senescent young-donor ASCs (i.e. those positive for senescence-associated (SA)-β-galactosidase) was 1.9 ± 0.8% at P3 and 22.2 ± 4.2% at P11 (Fig. 1c-d). Interestingly, these percentages were higher for aged-donor ASCs, with 6.0 ± 2.1% at P3 and 32.6 ± 4.1% at P11 (Fig. 1c-d). Accordingly, aged-donor ASCs displayed greater levels of lysosome accumulation (a hallmark of aging), as measured by Lysotracker fluorescence (Fig. 1e). The aged-donor ASCs also expressed significantly greater levels (vs. young-donor ASCs) of the cell cycle arrest markers p16^INK4^ and p21^WAF1^ and of the pro-senescence protein prelamin A from P3 onwards (Fig. 1f-g). These observations suggest that ASCs from aged donors presented signs of senescence earlier in culture than those derived from young donors.

**Figure 1:**
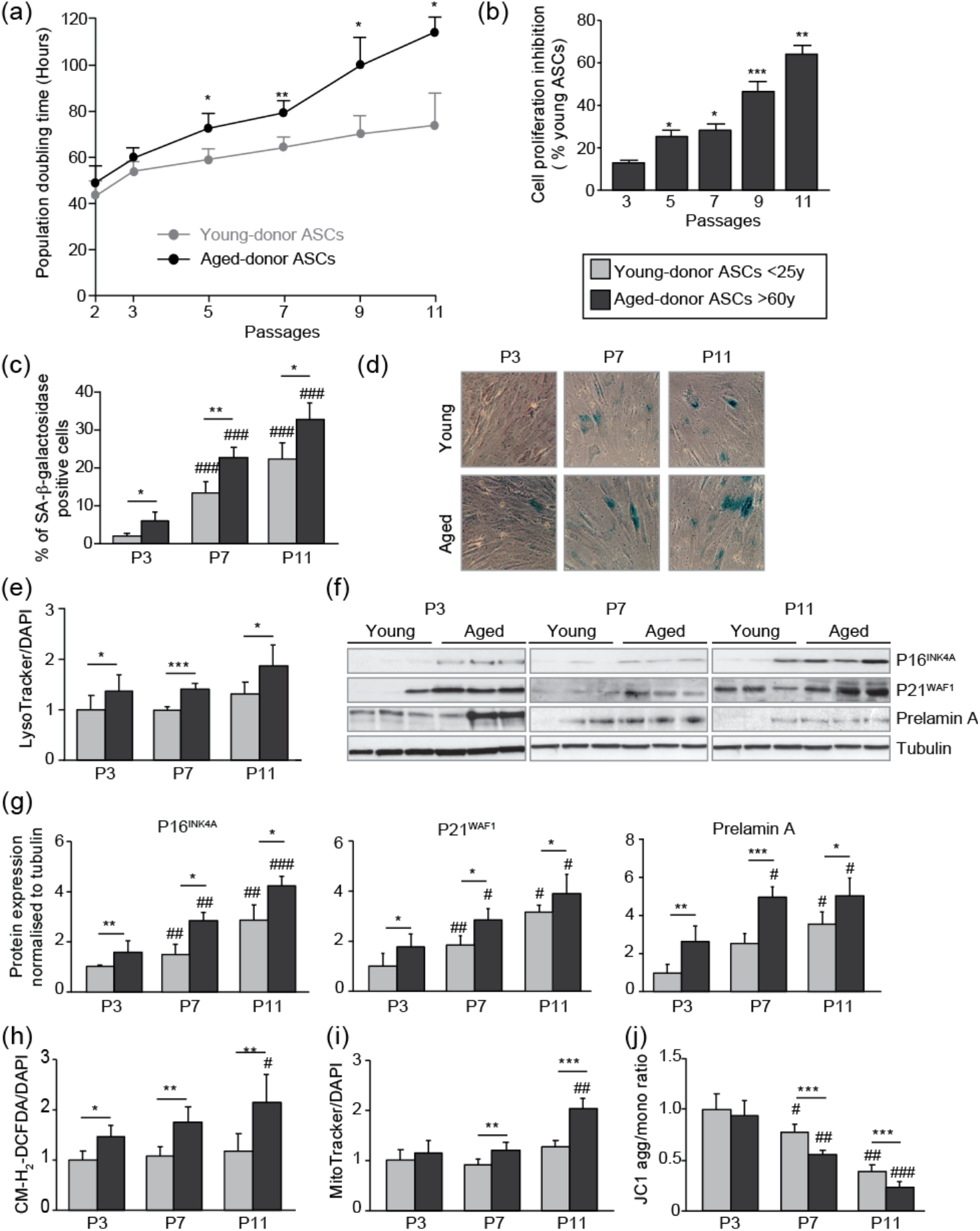
*In vivo* aging is associated with earlier *in vitro* senescence and greater oxidative stress and mitochondrial dysfunction in cultured ASCs. After isolation from the abdominal SCAT of young and aged donors, ASCs were cultured from P3 to P11. (**a**). Calculation of the mean ± SEM population doubling time (PDT) is described in the Material and Methods section. Times were determined at the indicated passage (n=9, in triplicate). (**b**) The % inhibition of cell proliferation was calculated for aged-donor ASCs by determining the increase in total cell number that occurred after 7 days, compared to young-donor ASCs. (**c**) Senescence was evaluated in terms of SA-β-galactosidase activity and expressed as the proportion (in %) of SA-β-galactosidase-positive cells at pH6 in aged-donor ASCs vs. young-donor ASCs at the same passage (at P3, P7 and P11). (**d**) Representative micrographs of SA-β-galactosidase-positive cells. (**e**) Lysosomal accumulation (normalized against DAPI) was assessed with the Lysotracker fluorescence probe and expressed as the fluorescence ratio for aged-donor ASCs vs. young-donor ASCs at P3. (**f**) Whole-cell lysates of aged-donor and young-donor ASCs at P3, P7 and P11 were analysed by immunoblotting. Representative immunoblots of the cell cycle arrest markers p16INK4A and p21WAF1, prelamin A, and tubulin (the loading control) are shown. (**g**) Quantification of western blot were normalized to young-donor ASCs at P3. (**h**) Reactive oxygen species production (normalized against DAPI) was assessed by the oxidation of CM-H2DCFDA and expressed as a ratio relative to young-donor ASCs at P3. (**i**) Mitochondrial mass (normalized against DAPI) was evaluated with Mitotracker Red-Probe and expressed as a ratio relative to young-donor ASCs at P3. (**j**) The cationic dye JC1 was used to evaluate the mitochondrial membrane potential. The results are expressed as the ratio of aggregate/monomer fluorescence. Results are quoted as the mean ± SEM. *P<0.05, **P<0.01, ***P<0.001 for aged-donor *vs*. young-donor ASCs, #P<0.05, ##P<0.01, ###P<0.001 *vs*. young- or aged-donor ASCs at P3. All experiments were performed in duplicate or triplicate with ASCs isolated from 5 different donors in each group.

Lastly, aged-donor ASCs produced more ROS that young-donor ASCs at all cell passages (Fig. 1 h). We used Mitotracker to label mitochondria and evaluated their volume (higher volume being a sign of mitochondrial dysfunction). Although young- and aged-donor ASCs had similar volumes at P3, this variable was significantly higher in aged-donor ASCs at P7 and especially at P11 (Fig. 1i). We also observed destabilisation of the mitochondrial membrane potential in aged-donor ASCs, as shown by the JC1 assay (Fig. 1j). The level of JC1 fluorescence in aged-donor cells had fallen by 29% at P7 and by 40% at P11 (relative to young-donor ASCs), in line with the time of onset of mitochondrial dysfunction.

Taken as a whole, these results show that aged-donor ASCs experienced senescence earlier and more intensely than young-donor ASCs did. The phenotypic differences were maintained throughout the culture period – suggesting that ASCs may recapitulate the *in vivo* aging process *in vitro*.

### Aging is associated with earlier dysfunction in adipocyte-differentiated ASCs, and alters the latter’s adipogenic differentiation capacity

We next evaluated the impact of donor age on the ASCs’ ability to differentiate into mature adipocytes. At passages 3, 7 or 11, confluent cells were induced to differentiate for 14 days in a pro-adipogenic medium. We observed a gradually decrease in staining with Oil-Red-O (a marker of triglyceride accumulation) between P3 and P11 in both young- and aged-donor adipocytes, suggesting that increasing passage negatively impacts adipogenesis (Fig. 2a-b). Moreover, aged-donor adipocytes displayed less lipid accumulation than young-donor adipocytes at P7 (21% less) and P11 (35% less). Accordingly, the protein expression level of the main adipogenic transcription factors C/EBPα, SREBP1c and PPARγ decreased with increasing passage and was even lower in aged-donor adipocytes than in young-donor adipocytes (Fig. 2c-d). Interestingly, aged-donor adipocytes did not display a delay in adipogenesis at P3 but their levels of oxidative stress (Fig. 2e) and mitochondrial dysfunction were already higher than in young-donor cells (Fig. 2f-g). These differences were accentuated at later passages; at P11, we observed 3-fold greater ROS production, 1.8-fold greater mitochondrial mass, and a 25% lower mitochondrial membrane potential.

**Figure 2:**
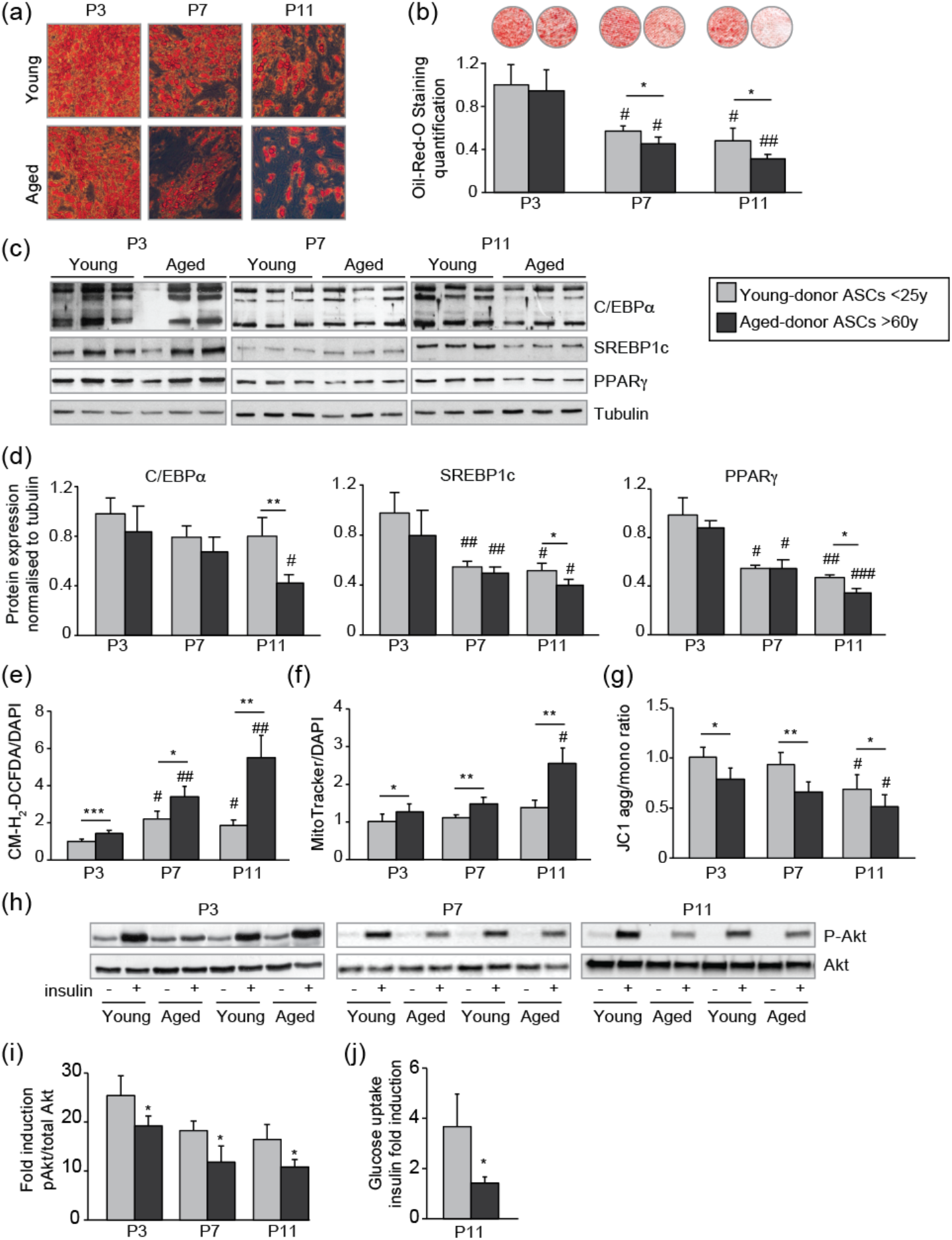
*In vivo* aging is associated with earlier *in vitro* dysfunction and altered adipogenesis of adipocytes differentiated from ASCs. The ASCs were differentiated into adipocytes for 14 days at P3, P7 and P11. **(a)** Cells were stained with Oil-Red-O to visualize lipid droplets 14 days post-induction, and representative micrographs are shown. **(b)** Quantification of Oil-Red-O staining of adipocytes differentiated from ASCs and representative scans of wells are shown. **(c)** Whole-cell lysates at day 14 post-induction of adipocytes differentiated from ASCs isolated from young and aged donors cultured until P3, P7 and P11 were analysed by immunoblotting. Representative immunoblots of C/EBPα, SREBP-1c, PPARγ and tubulin (loading control) are shown. **(d)** Quantification of western blot were normalized to young-donor ASCs at P3 **(e)** ROS production, normalized against DAPI. **(f)** Mitochondrial mass (normalized against DAPI) and **(g)** mitochondrial membrane potential were assessed in aged-donor ASCs as described in Figure 1. **(h)** Whole-cell lysates (extracted at day 14 post-induction) of adipocytes differentiated from young- and aged-donor ASCs and stimulated (or not) with insulin were analysed with immunoblotting. Representative immunoblots of Akt and phospho-Akt (Ser473) are shown. (**i**) The phosphorylated Akt/total Akt ratio was determined in a densitometric analysis. **(j)** Insulin sensitivity at P11 in adipocytes differentiated from young- and aged-donor ASCs was evaluated by measuring glucose uptake in basal and insulin-stimulated conditions as described in the Material and Methods section. The insulin fold induction was determined. Results are quoted as the mean ± SEM. *P<0.05, **P<0.01, ***P<0.001 for aged-donor *vs*. young-donor ASCs, #P<0.05, ##P<0.01, ###P<0.001 *vs*. young- or aged-donor ASCs at P3. All experiments were performed in duplicate or triplicate with ASCs isolated from 5 different donors in each group.

We also evaluated the insulin sensitivity of adipocytes derived from young- and aged-donor ASCs. A Western blot analysis showed that both groups of adipocytes expressed similar levels of Akt, a key enzyme involved in short-term metabolic responses to insulin (Fig. 2h). Nonetheless, aged-donor adipocytes displayed low phosphorylation of Akt in response to insulin at all passages (Fig. 2h-i) – suggesting that despite complete differentiation from ASCs at early passages, adipocytes differentiated from aged-donor ASCs were already insulin-resistant. At P11, aged-donor adipocytes showed a lower insulin-induced glucose transport (1.4 fold) response than young-donor cells (3.7 fold) (Fig. 2j).

### Metformin prevents the onset of senescence and associated dysfunctions in ASCs isolated from aged donors but not in ASCs from young donors

Next, we looked at whether metformin could alleviate cellular senescence in ASCs. To that end, ASCs were treated with metformin from P3 to P11. As shown in Figure 3a, metformin did not modify the PDT of young-donor ASCs. However, metformin prevented the above-mentioned relative decrease in cell proliferation in aged-donor ASCs (Fig. 3a-b). Accordingly, metformin rescued the higher percentage of senescent cells (Fig. 3c-d), the greater lysosome accumulation (Fig. 3e), and the higher expression of cell cycle inhibitors p21^WAF1^ and p16^INK4^ (Fig. 3f) previously observed at P11. Taken as a whole, these data show that in aged-donor ASCs, metformin reverted the senescence-associated dysfunction, oxidative stress and mitochondrial dysfunction to the levels observed in young-donor ASCs (Fig. 3g-i).

**Figure 3:**
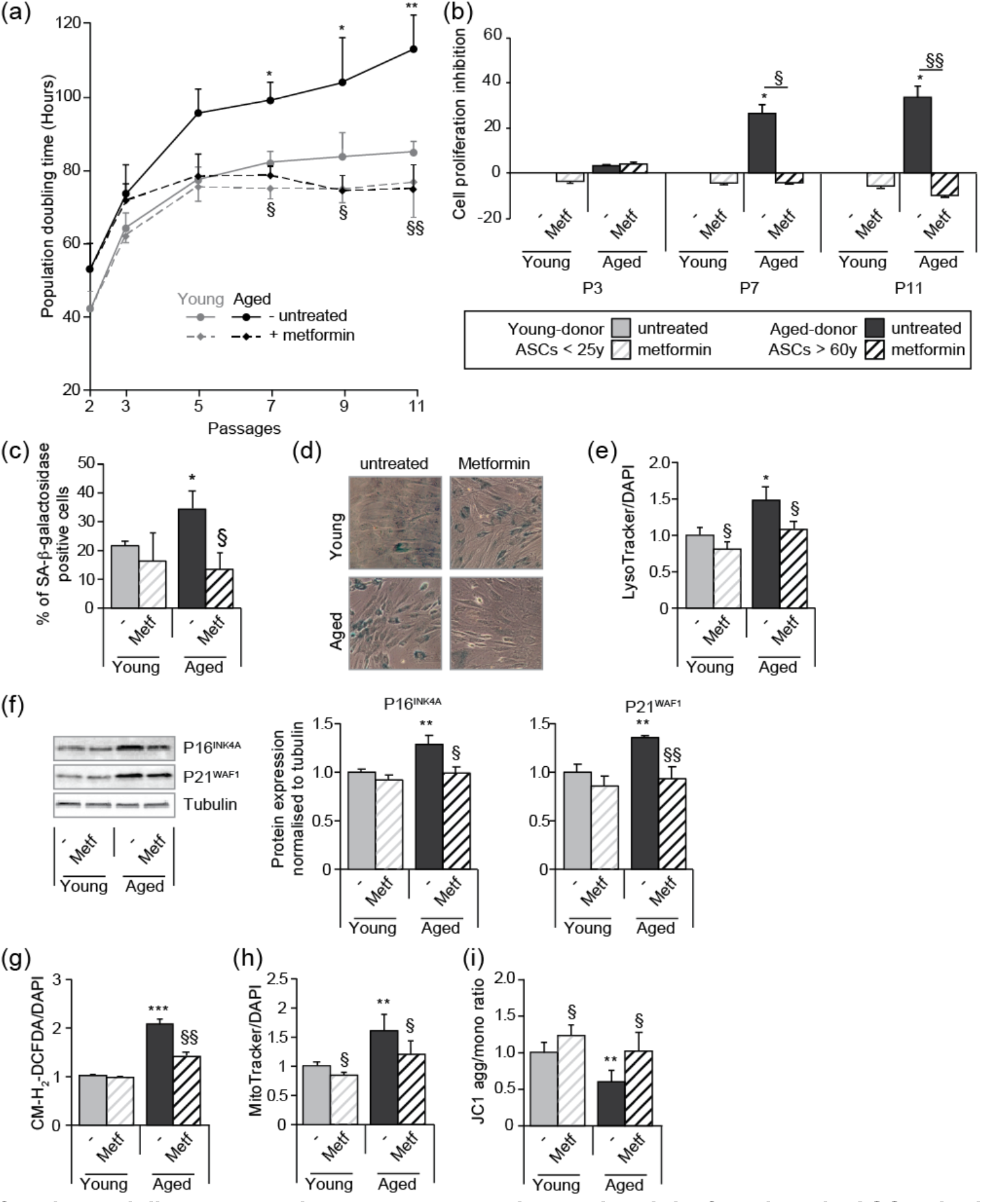
Metformin partially prevents the senescence and associated dysfunctions in ASCs obtained from aged donors but not in those obtained from young donors. Metformin (25 μM) was added to the culture medium of aged-donor and young-donor ASCs from P3 onwards. (**a)** Mean PDTs were determined at the indicated passages in aged-donor and young-donor ASCs treated (or not) with metformin at the same passage. (**b**) The % inhibition of cell proliferation was calculated for aged-donor ASCs and young-donor ASCs treated by metformin by determining the increase in total cell number that occurred after 7 days, compared to young-donor ASCs. (**c**) Senescence was evaluated in terms of SA-β-galactosidase activity and expressed as the proportion (in %) of SA-β-galactosidase-positive cells at pH6 in metformin-treated ASCs vs. non-treated ASCs at P11 (**d**) Representative micrographs of SA-β-galactosidase positive cells. (**e**) Lysosomal accumulation (normalized against DAPI) was assessed with the Lysotracker fluorescence probe in metformin-treated ASCs vs. non-treated ASCs at P11. (**f**) Whole-cell lysates of aged-donor and young-donor ASCs treated (or not) with metformin were analysed at P11 by immunoblotting. Representative immunoblots of the cell cycle arrest markers p16INK4A and p21WAF1 and of tubulin (the loading control) are shown. Quantitation of Western blots, normalized against the values for non-treated young-donor ASCs at P11. (**g**) ROS production, (**h**) mitochondrial mass (both normalized against DAPI) and (**i**) mitochondrial membrane potential were assessed as described in Figure 1 in metformin-treated ASCs *vs*. non-treated ASCs at P11. The results correspond to the mean ± SEM. *P<0.05, **P<0.01, ***P<0.001 for aged-donor *vs*. young-donor ASCs, §P<0.05, §§P<0.01 metformin-treated *vs*. non-treated ASCs. All experiments were performed in duplicate or triplicate in ASCs isolated from 3 different donors in each group.

### Metformin restores the ability of aged-donor ASCs to differentiate into adipocytes

We checked whether metformin pre-treatment was able to rescue the altered adipogenesis potential of aged-donor ASCs. Although the cells were treated with metformin, adipogenesis was induced in the absence of the drug to bypass its anti-adipogenic impact (Marycz, Tomaszewski et al. 2016, Chen, Wang et al. 2018). We found that metformin restored the adipogenic capacity of aged-donor ASCs at P11 to the level observed in young-donor ASCs. Indeed, metformin treatment during the proliferation state normalized lipid accumulation (Fig. 4a-b) and increased C/EBPα, and SREBP1c expression (Fig. 4c-d). Accordingly, metformin treatment accentuated the insulin response in adipocytes (Fig. 4e-f). Lastly, metformin pre-treatment lowered levels of oxidative stress (Fig. 4g) but did not restore mitochondrial function (Fig. 4h-i).

**Figure 4:**
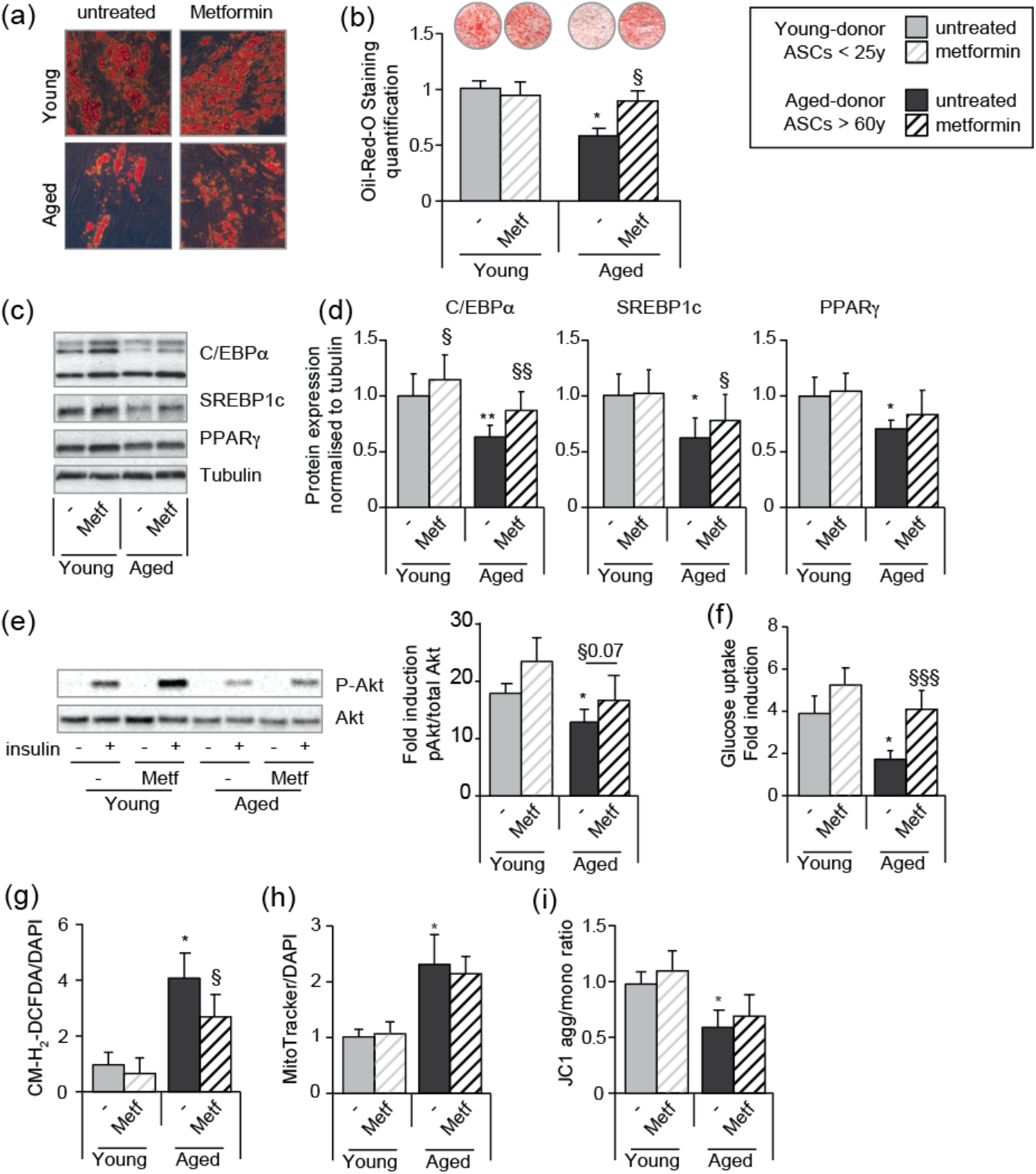
Metformin can restore the adipogenic differentiation capacity of senescent aged-donor ASCs. Metformin was added to the culture medium of young- and aged-donor ASCs from P3 onwards. The ASCs were differentiated into adipocytes for 14 days at P11, in the absence of metformin. **(a)** Cells were stained with Oil-Red-O to visualize lipid droplets 14 days post-induction, and representative micrographs are shown. **(b)** Quantification of Oil-Red-O staining and representative scans of wells are shown. **(c)** Whole-cell lysates on day 14 post-induction from adipocytes differentiated from non-treated young-donor and aged-donor ASCs at P11 and treated (or not) with metformin were analysed by immunoblotting. Representative immunoblots of C/EBPα, SREBP-1c, PPARγ and tubulin (the loading control) are shown. **(d)** Quantification of western blots is shown. **(e)** Whole-cell lysates extracted at day 14 post-induction stimulated (or not) by insulin from differentiated ASCs, were analysed by immunoblotting. Representative immunoblots of Akt and phospho-Akt (Ser473) and quantification of the pAkt/Akt are shown. **(f)** Insulin sensitivity was evaluated at P11 in adipocytes differentiated from non-treated young-donor and aged-donor ASCs treated (or not) with metformin, by measuring the glucose uptake in response to insulin and calculating the insulin fold induction, as described in the Material and Methods section. **(g)** ROS production, **(h)** mitochondrial mass and **(i)** mitochondrial membrane potential (both normalized against DAPI) were assessed as described in Figure 1. The results are expressed as the mean ± SEM. *P<0.05 for adipocytes differentiated from aged-donor *vs*. adipocytes differentiated from young-donor ASCs, §P<0.05, §§P<0.01, §§§P<0.001 metformin-treated *vs*. non-treated adipocytes differentiated from ASCs. All experiments were performed in duplicate on cells isolated from 3 different donors in each group.

### The beneficial effect of metformin on ASC senescence is mediated by AMPK activation

We hypothesized that in mechanistic terms, metformin’s action might be based on (amongst other things) the phosphorylation and thus activation of AMPK. As shown in Figure 5a, metformin treatment was associated with (i) greater AMPK expression in both young-donor and aged-donor ASCs but (ii) greater AMPK phosphorylation in aged-donor ASCs only. These findings are in line with metformin’s beneficial effect on aged-donor ASCs and lack of effect on young-donor ASCs, as reported above.

**Figure 5:**
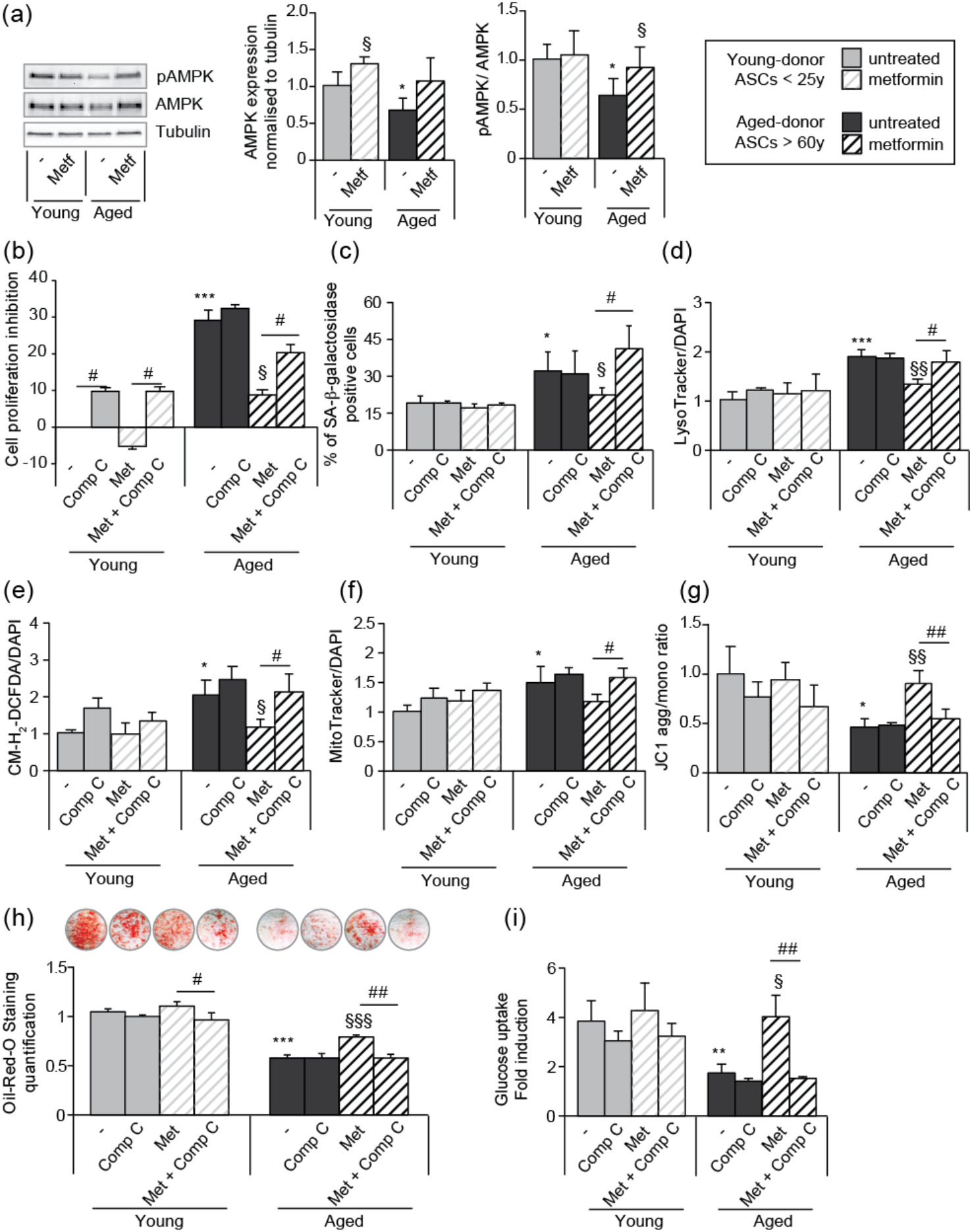
The beneficial effects of metformin on aged-donor ASC senescence is mediated by AMPK activation. Metformin was added to the culture medium of aged-donor and young-donor ASCs from P3 onwards. To evaluate the role of AMPK activation, compound C was added at P11. After 7 days of treatment, the experiments on ASCs were carried out. **(a)** Whole-cell lysates of aged-donor and young-donor ASCs treated (or not) with metformin and compound C at P11 were analysed by immunoblotting. Representative immunoblots of AMPK, phospho-AMPK and tubulin (the loading control) and a graph quantifying AMPK (normalized against tubulin) and the pAMPK/AMPK ratio are shown**. (b)** The % inhibition of cell proliferation was calculated for aged-donor ASCs and young-donor ASC treated or not with metformin or compound C, by determining the increase in total cell number that occurred after 7 days, compared to young-donor ASCs. **(c)** Senescence was evaluated in terms of SA-β-galactosidase activity and was expressed as described in Figure 1. **(d)** Lysosomal accumulation (normalized against DAPI) was assessed with the Lysotracker fluorescence probe. **(e)** ROS production, **(f)** mitochondrial mass (both normalized against DAPI) and **(g)** mitochondrial membrane potential were assessed as described in Figure 1. **(h)** The ASCs were then differentiated into adipocytes on P11 in the absence of metformin and compound C. Cells were stained with Oil-Red-O to visualize lipid droplets 14 days post-induction. Quantification of Oil-Red-O staining and representative scans of wells are shown. **(i)** Insulin sensitivity was evaluated at P11 in adipocytes differentiated from non-treated young-donor and aged-donor ASCs treated (or not) with metformin, by measuring the glucose uptake in response to insulin and calculating the insulin fold induction, as described in the Material and Methods section. Results are expressed as the mean ± SEM. *P<0.05, **P<0.01 for aged-donor *vs*. young-donor ASCs, §P<0.05, §§P<0.01 for metformin-treated ASCs *vs*. non-treated ASCs. #P<0.05, ##P<0.01 for compound C-treated ASCs *vs*. non-treated or metformin-treated ASCs. All experiments were performed in duplicate or triplicate on ASCs isolated from at 3 different donors in each group.

To confirm AMPK’s potential role in the action of metformin, we determined whether the drug’s effects were influenced by treatment with the AMPK inhibitor compound C. Indeed, we observed that in aged-donor ASCs, treatment with compound C counteracted the beneficial effect of metformin on cell proliferation (Fig. 5b), senescence marker levels (Fig. 5c-d), oxidative stress (Fig. 5e), and mitochondrial dysfunction (Fig. 5f-g). Interestingly, adipocytes derived from aged-donor ASCs and then pre-treated with both metformin and compound C during proliferation but not during differentiation did not show metformin’s beneficial effect on adipogenesis (fig. 5h) and insulin-stimulated glucose uptake (Fig. 5i). These findings highlighted AMPK’s role in the beneficial action of metformin on aged-donor ASCs and the derived adipocytes.

## DISCUSSION

Our present results show that *in vitro* culture reconstitutes some of the aging-associated changes that have been observed in ASCs *in vivo*. Indeed, long-term cultured *in vitro* ASCs presented the main characteristics of senescent cells. Moreover, when ASCs were obtained from aged donors, all the aging features were enhanced as compared to young-donor ASCs as a result of greater senescence. Thus, the aged-donor ASCs’ proliferative capacity was reduced with higher levels of several senescence biomarkers (including SA-β-galactosidase activity and cell cycle arrest proteins p21^WAF1^ and p16^INK4^) and with greater mitochondrial dysfunction and ROS production. In adipocytes differentiated from aged-donor ASCs, adipogenesis and insulin sensitivity were decreased as compared to young-donor ASCs. By activating AMPK, metformin restored almost all the features of aging in aged-donor ASCs to the levels observed in young-donor ASCs. However, metformin did not modify these variables in the latter cells.

Our data are in line with previous studies of human ASCs (Zhu, Kohan et al. 2009, Jung, Volk et al. 2019, Alicka, Kornicka-Garbowska et al. 2020). The ASCs have a typical mesenchymal stem cell morphology, and there are little or no morphological differences between young-donor and aged-donor cells in early passages. In most studies, the PDT of human ASCs did not vary with age (Zhu, Kohan et al. 2009, Chen, Lee et al. 2012, Ding, Chou et al. 2013). Accordingly, we did not observe any differences between young-donor and aged-donor ASCs with regard to the phenotype or proliferation rate early in the period of culture (P3). However, with increasing passage numbers, aged-donor cells developed features of senescence (such as increased SA-β-galactosidase activity, oxidative stress, and mitochondrial dysfunctions) earlier than young-donor cells. These alterations might have driven the progressive fall in proliferation capacity seen for aged-donor ASC from P5 onwards. Indeed, we found that aged-donor ASCs had molecular, morphological and functional impairments late in culture (P11). We propose that our results provide a better understanding of the molecular events occurring in ASCs in the course of aging; they recapitulated the *in vivo* aging of ASCs and revealed the peculiar features associated with the aging process in older compared to younger individuals.

With regard to the mechanisms that might promote senescence in ASCs, greater mitochondrial dysfunction and ROS production have previously been linked to the occurrence of age-dependent ASC senescence. Poor mitochondrial function and elevated mitochondrial generation of ROS are thought to be critical in the aging process (Seo, Joseph et al. 2010, Correia-Melo, Marques et al. 2016). Moreover, ROS accumulation was observed at P3 whereas mitochondrial dysfunctions were only observed at P7 suggesting that ROS could initially not originate from mitochondria.

We also looked at the impact of aging on the adipogenic fate of ASCs. Although the ASCs did not display delayed adipogenesis at P3, the presence of oxidative stress and mitochondrial dysfunction was a harbinger of impaired adipogenesis. Indeed, the ability to differentiate into adipocytes at P11 was significantly decreased in aged-donor ASCs. This finding is consistent with several other studies of human and murine mesenchymal stem cells having shown a reduction in differentiation potential upon aging *in vitro* (Choudhery, Badowski et al. 2014, Maredziak, Marycz et al. 2016). Moreover, we identified the early onset (at P3) of cellular insulin resistance in aged-donor ASCs for the first time that could participate to impaired adipogenesis.

One of the study’s objectives was to determine whether preventing the onset of senescence in aged-donor ASCs restored their adipogenic capacity. There is experimental evidence that metformin extends lifespan in model organisms, sparking interest in its clinical relevance (Martin-Montalvo, Mercken et al. 2013). We used a metformin concentration of 25 μM, in the upper range observed in diabetic patients and considered as a safe concentration (Frid, Sterner et al. 2010). Previous studies have demonstrated that metformin significantly improved ASC proliferation and function, which were correlated with a higher mitochondrial membrane potential (Smieszek, Kornicka et al. 2019). Metformin was shown to decrease ROS production by ASCs (Marycz, Tomaszewski et al. 2016). Here, we showed that aged-donor ASCs expressed and activated AMPK to a lower extent than young-donor ASCs, and that this impairment could be reversed by metformin treatment. This effect is related to metformin’s ability to inhibit complex 1 in the mitochondrial electron transport chain, leading to activation of AMPK and a reduction in endogenous ROS production. Accordingly, we found that metformin was a potent antioxidant and prevented the onset of mitochondrial dysfunction in aged-donor ASCs. Interestingly, we found that metformin treatment had a beneficial effect on aged-donor ASCs but not on young-donor ASCs; this suggests that *in vitro* and *in vivo* aging have different mechanisms, even though they result in similar cellular and molecular alterations. It has been shown that metformin has various effects on several pathways and targets, in addition to AMPK (Barzilai, Crandall et al. 2016). Even though metformin’s ability to inhibit mitochondrial complex 1 has been best characterized, many other pathways are affected and metformin’s mechanisms of action still warrant further investigation. Nonetheless, we observed that beneficial effects of metformin were lost in the presence of compound C (a potent AMPK inhibitor) – highlighting the central role of the AMPK-linked signalling network in the ASC aging process (Salminen and Kaarniranta 2012, Burkewitz, Zhang et al. 2014)

To the best of our knowledge, the present study is the first to have shown that metformin can partially rescue/maintain the adipogenic capacity of aged-donor ASCs. Metformin has been shown to inhibit adipogenesis (Marycz, Tomaszewski et al. 2016, Chen, Wang et al. 2018) when cells are exposed during the process of differentiation. Here, we used a novel model in which ASCs were only treated with metformin before the induction of differentiation. By removing metformin during ASC differentiation, we were able to rule out a direct effect on adipogenesis. Thus, our data suggest that metformin’s prevention of senescence and dysfunction during ASC proliferation restored adipogenesis. Interestingly, we also observed that metformin increased the insulin sensitivity of newly differentiated ASCs issued from aged-donor ASCs by activating AMPK; this activation has been shown to contribute to the drug’s insulin-sensitizing effect *in vivo* – particularly in the liver and in muscle (Salminen and Kaarniranta 2012).

The present study had several limitations. Firstly, we used ASCs obtained from abdominal SCAT; ASCs obtained from other subcutaneous or VAT depots might not behave in the same way. We found that ASCs are affected both by chronological *in vivo* aging and *in vitro* aging induced by longterm culture. It remains to be established whether these two processes are different. It has previously been suggested that both *in vivo* and *in vitro* aging of human ASCs induce similar alterations in gene expression (Wagner, Bork et al. 2009, Geissler, Textor et al. 2012). Another limitation concerns the use of fibroblast growth factor 2 (FGF2), which is required for the prolonged *in vitro* expansion of ASCs. It has been shown that FGF2-treated ASCs are larger, less readily proliferative, and more senescent than control ASCs (Cheng, Lin et al. 2020). Thus, we cannot rule out a possible masking effect of FGF2 on the aging of ASCs.

Aging is associated with the induction of senescence and associated disorders (including oxidative stress, and insulin resistance) in ASCs. These changes might be involved in the alterations in fat redistribution (particularly a paucity of SCAT, favouring VAT accumulation) and cardiometabolic diseases observed during aging. Our results suggest that metformin may be a promising candidate for treating age-related dysfunction in adipose tissue (Fig. 6), and thus warrant evaluation in a clinical setting.

**Figure 6:**
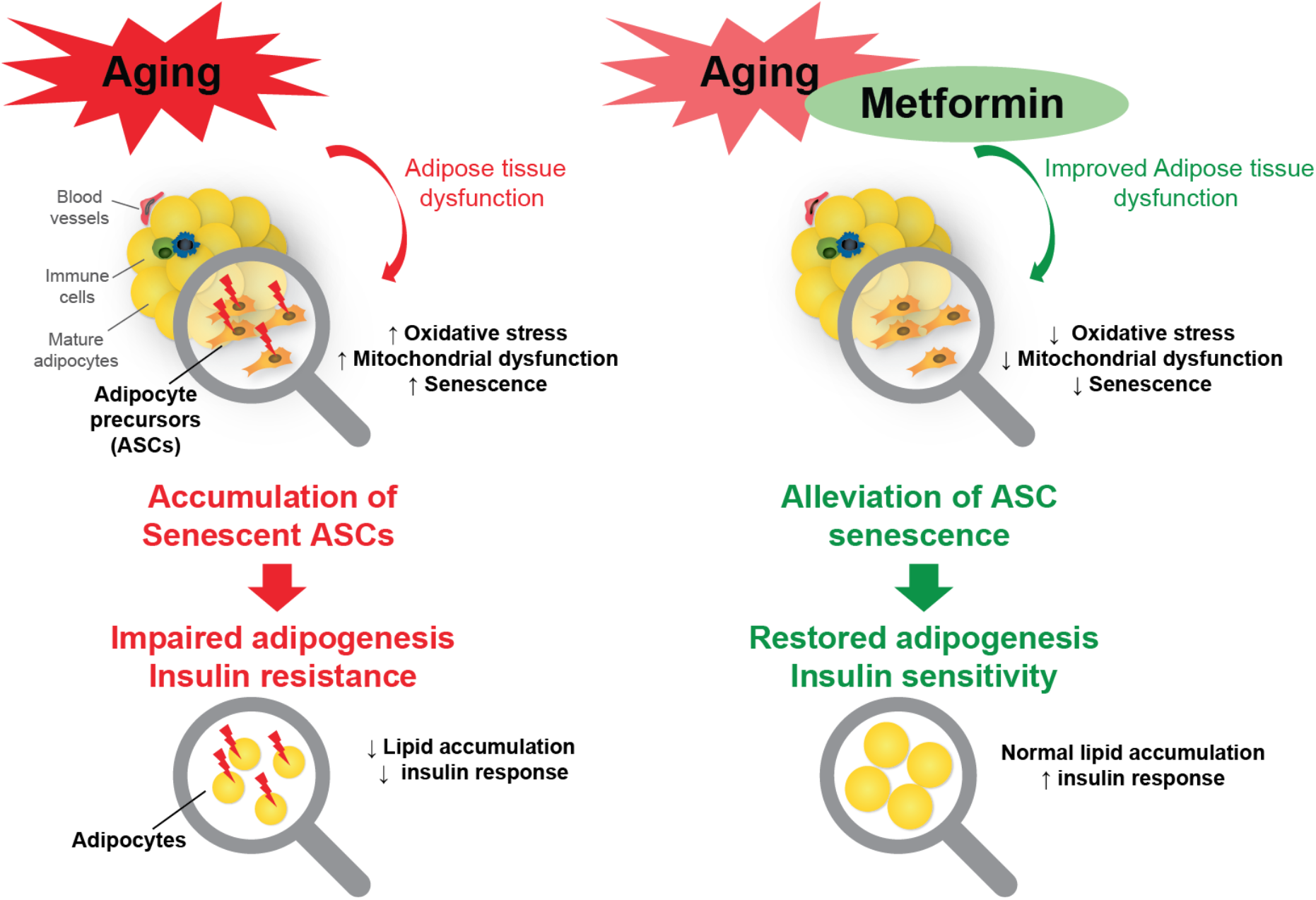
Targeting human adipose stem cell senescence with metformin enhances adipogenesis and insulin sensitivity.

## MATERIAL AND METHODS

### Isolation, culture and treatment of ASCs

The human SCAT samples from which ASCs were isolated were obtained from 10 healthy women undergoing plastic surgery. The women were young adults (n=5; mean ± standard error of the mean (SEM) age: 21.2 ± 3.2; mean ± SEM BMI: 23.6 ± 1.2 kg/m^2^) or aged adults (n=5; mean ± SEM age: 60.0 ± 0.9; mean ± SEM BMI: 25.7 ± 0.5 kg/m^2^). Before surgery, all donors provided informed written consent to use of their tissue specimens for research purposes. The study was performed in compliance with the principles of the Declaration of Helsinki, and was approved by the local independent ethics committee. The ASCs were isolated using a collagenase (Roche Diagnostics, Basel, Switzerland) digestion technique, as described previously (Gorwood, Bourgeois et al. 2019, Gorwood, Ejlalmanesh et al. 2020), after seeding in Eagle’s Minimum Essential Medium alpha (Thermo Fisher Scientific, Courtaboeuf, France) supplemented with 10% foetal bovine serum (FBS) (PAN-Biotech, Aidenbach, Germany), 2 mmol/L glutamine, 100 U/mL penicillin/streptomycin, 10 mmol/L HEPES (Thermo Fisher Scientific), and 2.5 ng/mL FGF2 (PeproTech, Rocky Hill, NJ, USA). Upon confluence, adherent cells were trypsinized (Thermo Fisher Scientific) and seeded at a density of between 2000 and 4000 cells/cm^2^. During expansion, cells were exposed (or not) to 25 μmol/L metformin (Merck, Sigma Aldrich, St. Quentin Fallavier, France). The metformin concentration chosen in our study is close to Cmax observed in ederly subjects, which exhibited higher average Cmax than younger subjects (Jang, Chung et al. 2016). At a late passage (P11), cells were exposed (or not) to 0.1 μmol/L compound C (Merck, Sigma Aldrich).

### Adipocyte differentiation

Differentiation of ASCs was induced by the addition of proadipogenic Dulbecco Modified Eagle’s Medium (DMEM), 4.5 g/L glucose (Thermo Fisher Scientific), 10% FBS, 2 mmol/L glutamine, 100 U/mL penicillin/streptomycin, 10 mmol/L HEPES, 1 μmol/L dexamethasone, 250 μmol/L 3-Isobutyl-1-methylxanthine (IBMX), 1 μmol/L rosiglitazone, 1 μmol/L insulin (Merck, Sigma-Aldrich) for 5 days, and then maintained in DMEM with rosiglitazone and insulin up until day 14. Cells were then stained for Oil-Red-O (Merck, Sigma-Aldrich) and quantified at 520nm as described previously (Gorwood, Bourgeois et al. 2019, Gorwood, Ejlalmanesh et al. 2020).

### Cellular proliferation and senescence

Cellular senescence was evaluated in terms of the cell PDT at each cell passage, as described previously (Gorwood, Ejlalmanesh et al. 2020). The positive blue staining of senescence-associated SA-β-galactosidase has been used as a biomarker of cellular senescence. To detect SA-β-galactosidase activity, cells were incubated in an appropriate buffer solution at pH6 containing bromo-4-chloro-3-indolyl-β-D-galactopyranoside (Euromedex, Souffelweyersheim, France), as described previously (Gorwood, Ejlalmanesh et al. 2020). The percentage of blue SA-β-galactosidase-positive cells was estimated by cell counting in at least 3 random-selected fields at a magnification of 20X. The acidotropic dye LysoTracker (Invitrogen Corporation, Carlsbad, CA, USA) was used to evaluate the lysosomal mass. Cells cultured in 96-well plates (Corning, New York, NY USA) were incubated with Lysotracker for 2 h at 37°C. The fluorescence was quantified on a plate reader at 504 nm-570 nm (Tecan, Trappes, France), and normalized against 4’,6-Diamidino-2-phenylindole dihydrochloride (DAPI) fluorescence at 345 nm-458 nm.

### Mitochondrial dysfunctions and oxidative stress

The cationic dye tetra-chloro-tetra-ethyl-benzimidazolyl-carbocyanine iodide (JC1) was used to evaluate the mitochondrial membrane potential, and the Mitotracker Red probe (both from Invitrogen Corporation) was used to measure the mitochondrial mass. The production of ROS was assessed by the oxidation of 5-6-chloromethyl-2,7-dichlorodihydrofluorescein diacetate (CM-H_2_DCFDA) (Invitrogen Corporation). Cells cultured in 96-well plates were incubated with JC1, Mitotracker, or CM-H_2_DCFDA for 2 h at 37°C. The fluorescence was quantified on a plate reader at 520 nm-595 nm for JC1 aggregates, 485 nm-535 nm for JC1 monomers, 575 nm-620 nm for Mitotracker, and 485 nm-520 nm for CM-H_2_DCFDA. The results were normalized against DAPI fluorescence.

### Protein extraction and Western blotting

Proteins were extracted from cell monolayers as described previously (Gorwood, Bourgeois et al. 2019, Gorwood, Ejlalmanesh et al. 2020). After SDS-PAGE, the proteins were transferred to nitrocellulose membranes. Specific proteins were detected using antibodies against p16^INK4A^, p21^WAF1^ (BD Pharmingen, Franklin Lakes, NJ, USA), prelamin A, PPARγ, CEBPα, SREBP1c, AMPK, phospho-AMPK (SCBT, Dallas, TX, USA), and the protein loading control tubulin (Merck, Sigma-Aldrich). Immuno-reactive complexes were detected using HRP-conjugated secondary antibodies (Cell Signaling Technology, Danvers, MA, USA) and enhanced chemiluminescence (Thermo Fisher Scientific).

### Insulin signalling and glucose transport

The insulin sensitivity of adipocyte-derived ASCs at late passage (P11) was evaluated by the phosphorylation of Akt. On day 14, the adipocytes were serum-starved for 18 h and stimulated for 7 minutes with 100 nmol/L insulin. Cell lysates were immunoblotted with antibodies against the activated forms (Ser473 phosphorylation) of Akt (Cell Signaling Technology). Protein expression was checked using antibodies against Akt (Cell Signaling Technology). Insulin-stimulated glucose uptake was measured using the Glucose Uptake-Glo assay kit (Promega, Fitchburg, WI, USA), according to the manufacturer’s instructions. Briefly, on day 14, adipocytes were serum-starved for 18 hours. Prior to the experiment, cells were incubated for 1 h in glucose-free DMEM (Thermo Fisher Scientific). Next, 100 nmol/L insulin and 2-deoxyglucose mix were successively added for 60 and 10 minutes, respectively. In parallel with the insulin-stimulated and non-stimulated conditions, 50 μmol/L of the actin-disrupting agent cytochalasin B (Merck, Sigma-Aldrich) was added as a negative control and enabled determination of the net insulin-stimulated glucose uptake. The reaction was stopped with neutralization buffer and detection reagent mix was added and incubated for 1 h at room temperature prior to measurement of luminescence on a plate reader.

### Statistical analysis

All experiments were performed in duplicate or triplicate on ASCs isolated from at least 3 or 5 different donors from each age group. All data were expressed as the mean ± SEM. The statistical significance of intergroup differences or changes over time was determined by applying a non-parametric test (the Mann–Whitney test), as appropriate.

## ACKNOWLEDGMENTS

We thank the patients for their cooperation. This research was funded by RHU CARMMA (grant number RHU-ANR-15-RHUS-0003), Fondation pour la Recherche Médicale (FRM, grant number EQU201903007868 to B.F.), the Institut National de la Santé et de la Recherche Médicale (INSERM), and Sorbonne Université.

